# Structure of the ciliary tip central pair reveals the unique role of the microtubule-seam binding protein SPEF1

**DOI:** 10.1101/2024.12.02.626492

**Authors:** Thibault Legal, Ewa Joachimiak, Mireya Parra, Wang Peng, Amanda Tam, Corbin Black, Melissa Valente-Paterno, Gary Brouhard, Jacek Gaertig, Dorota Wloga, Khanh Huy Bui

**Affiliations:** Department of Anatomy and Cell Biology, McGill University, Montreal, Québec, Canada; Laboratory of Cytoskeleton and Cilia Biology, Nencki Institute of Experimental Biology of Polish Academy of Sciences, 3 Pasteur Str, 02-093 Warsaw, Poland; Department of Cellular Biology, University of Georgia, Athens, GA, USA; Department of Biology, McGill University, Montreal, Québec, Canada; Centre de Recherche en Biologie Structurale, McGill University, Montreal, Québec, Canada

**Author notes:** Corresponding authors: Dorota Włoga, Khanh Huy Bui and. These authors contributed equally (Thibault Legal, Ewa Joachimiak).

**Keywords:** ciliary tip, central pair, axoneme, cryo-electron microscopy, microtubule

## Abstract

Motile cilia are unique organelles with the ability to autonomously move. Force generated by beating cilia propels cells and moves fluids. The ciliary skeleton is made of peripheral doublet microtubules and a central pair (CP) with a distinct structure at the tip. In this study, we present a high-resolution structure of the CP in the ciliary tip of the ciliate *Tetrahymena thermophila* and identify several tip proteins that bind and form unique patterns on both microtubules of the tip CP. Two of those proteins that contain tubulin polymerization-promoting protein (TPPP)-like domains, TLP1 and TLP2, bind to high curvature regions of the microtubule. TLP2, which contains two TPPP-like domains, is an unusually long protein that wraps laterally around half a microtubule and forms the bridge between the two microtubules. Moreover, we found that the conserved protein SPEF1 binds to both microtubule seams. *In vitro*, human SPEF1 not only binds to the microtubule seam but also crosslinks two parallel microtubules. Single-molecule microtubule dynamics assays indicate that SPEF1 stabilizes microtubules *in vitro*. Together, these data show that the proteins in the tip CP maintain stable microtubule structure and probably play important roles in maintaining the integrity of the axoneme.

## Introduction

Cilia are motile or sensory organelles found on the surface of many eukaryotic cells. Primary cilia are signalling hubs with fundamental roles in organism development, whereas motile cilia motility shifts fluids against ciliated surfaces, e.g., mucus flow in the airways. The main component of a cilium is a microtubular axoneme that is conserved across species (Klena and Pigino 2022). In motile cilia, the nine doublet microtubules surround the centrally positioned pair of microtubules (central pair, CP), forming the canonical 9+2 structure (Fig. 1A). Axonemal dyneins docked to the doublet microtubules generate the pulling forces required for ciliary beating, whereas the CP plays a crucial role in regulating ciliary motility through mechano-regulation (Klena and Pigino 2022).

**Figure 1:**
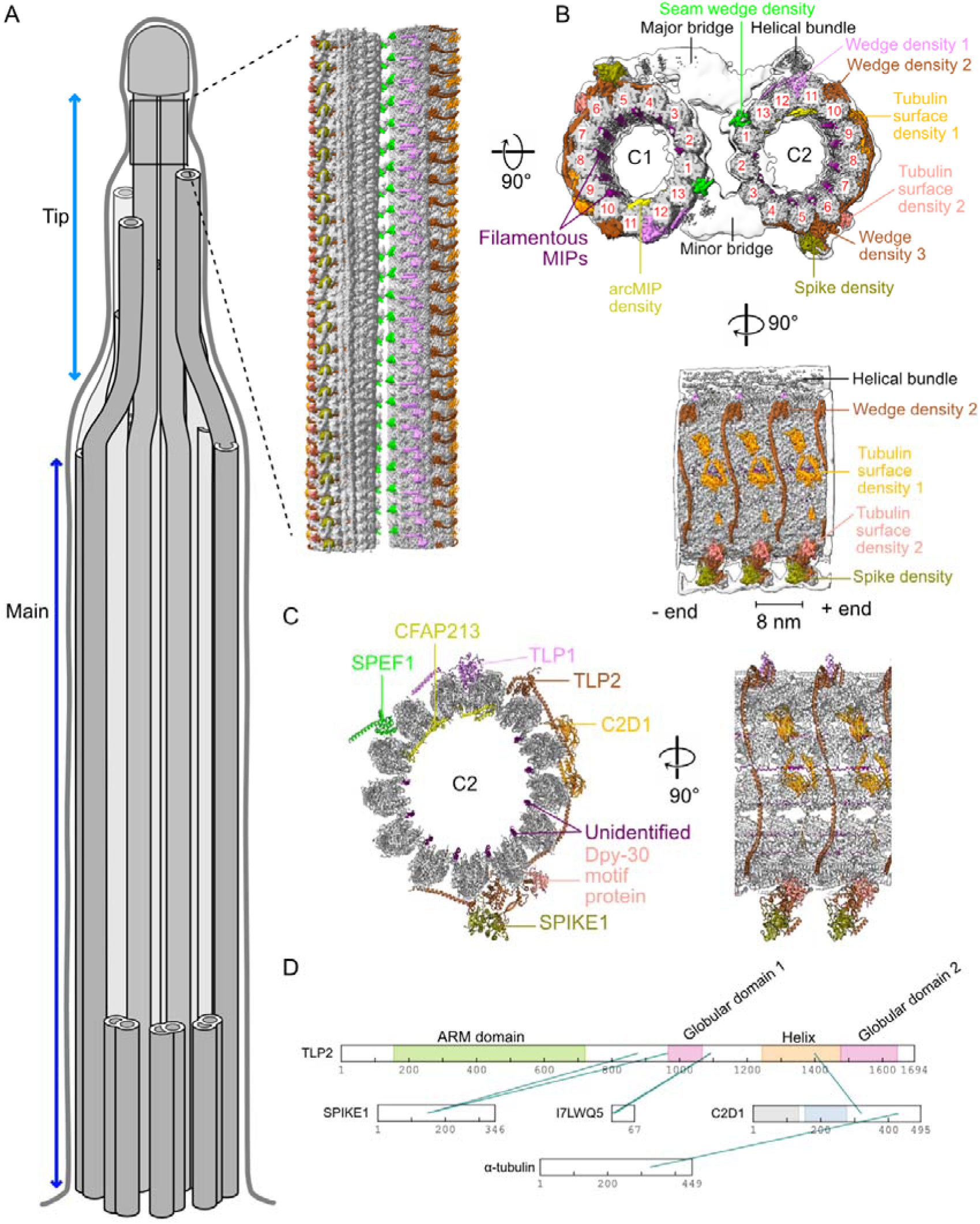
Atomic model of the tip CP. (A) Cartoon of cilia in Tetrahymena indicating the main and tip regions of the axoneme. The inset shows the cryo-EM map of the distinct structure of the tip CP. (B) Cryo-EM map of the tip CP. Tubulin is shown in gray, and the densities corresponding to the proteins identified in this study are shown in color. The low-resolution map of the tip CP (consensus map) is shown as an envelope in white. The PF numbers are shown in red. (C) Atomic model of C2 in the tip CP. Newly identified proteins that could be modelled are labelled in color. (D) In situ crosslinks identified among tip CP proteins and tubulins.

**Supplementary Figure 1:**
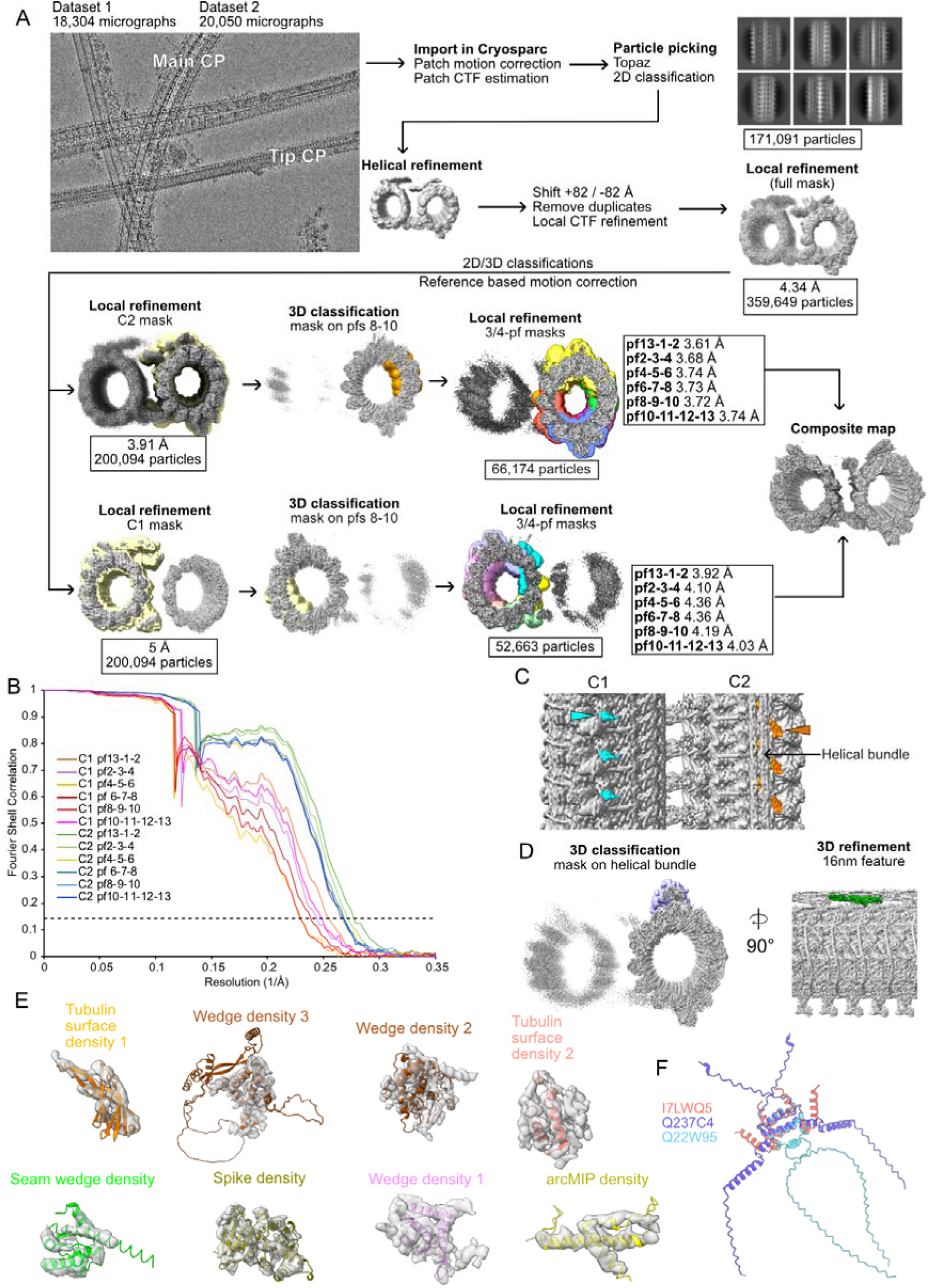
(A) Cryo-EM processing workflow of the tip CP. (B) Fourier shell correlation curves of the local refinements used in the composite map of the tip CP. (C) Densities specific to C1 (blue) and C2 (orange) highlight the asymmetry between the two microtubules. (D) Mask (purple) used for the 3D classification of the helical bundle and the 16 nm feature (green) identified. (E) Densities segmented for use with DomainFit (grey) and best fit of the predicted model for each density (colour). (F) AlphaFold2-predicted dimers of three ciliary DPY-30-containing proteins (UniProt IDs: I7LWQ5, Q237C4, and Q22W95).

While the 9+2 structure is maintained along the length of the cilium, the distal tip is a diverse and plastic structure (Soares, Carmona et al. 2019). The doublet microtubules transition into singlet microtubules in the tip region. In some species, the CP extends further than the singlet microtubules and is crowned by a cap complex (Dentler and Rosenbaum 1977, Suprenant and Dentler 1988). On the other hand, in sperm cells, the CP terminates at the same level or earlier than the outer singlets do (Fawcett 1965). In *Tetrahymena*, the CP extends approximately 500 nm in the tip region from the transition point between the doublet and singlet microtubules (Suprenant and Dentler 1988). A recent cryo-ET study of the ciliary tip in *Tetrahymena* indicated that the tip CP has an 8-nm repeating structure, which is markedly different from the main CP structure in the proximal and medial regions (Fig. 1A) (Legal, Parra et al. 2023).

There is limited understanding of the function of the tip region of the cilia, which is related mainly to its significant substoichiometric length. At the tip, microtubules interact with microtubule end-binding proteins and with intraflagellar transport motor proteins for assembly and maintenance (Soares, Carmona et al. 2019). Only a few ciliary tip proteins have been shown to localize to the tips of human primary cilia (Latour, Van De Weghe et al. 2020) or motile cilia (Louka, Vasudevan et al. 2018). Mutations in some proteins identified in the cilia tip cause Joubert syndrome, a neurodevelopmental disorder in humans, and severe ciliopathy phenotypes in zebrafish (Srour, Hamdan et al. 2015, Latour, Van De Weghe et al. 2020). SPEF1/CLAMP, a protein originally identified in mouse testes and characterized by the presence of the calponin homology (CH) domain (Chan, Fowler et al. 2005), localises along the length of cilia in mouse tracheal epithelial cells but accumulates at the tips of *Xenopus* epidermal multiciliated cells (Gray, Abitua et al. 2009, Zheng, Liu et al. 2019). SPEF1 is also expressed outside the cilia in large and organized microtubule bundles in mouse pillar cells in the organ of Corti in the cochlea (Dougherty, Adler et al. 2005). SPEF1 is essential for CP assembly (Zheng, Liu et al. 2019, Konjikusic, Lee et al. 2023). When overexpressed in fibroblasts, SPEF1 associates with microtubules, stabilizes them, and promotes bundle formation(Dougherty, Adler et al. 2005) (Dougherty, Adler et al. 2005). SPEF1 is also essential for central pair (CP) assembly (Zheng, Liu et al. 2019, Konjikusic, Lee et al. 2023). However, the ultrastructural location of SPEF1 within cilia and the molecular mechanism of its function are unknown.

Determination of the structure of the ciliary tip and identification of its specific proteins would enhance our understanding of microtubule plasticity in this region and suggest functions of tip-associated proteins.

In this study, we utilized cryo-electron microscopy (cryo-EM) to reconstruct the native tip-CP structure of *Tetrahymena*, only approximately one-sixtith of the total axonemal filament length. Through AlphaFold model prediction and domain fitting, we identified and modelled seven previously unknown tip proteins. Fluorescence tagging confirmed that six of these proteins localize specifically to the ciliary tip. The tip CP structure reveals distinct features of these proteins, including the presence of tubulin polymerization promoting protein (TPPP)-like domains in two of them, which preferentially associate with regions of high microtubule curvature. TLP2, a TPPP-like domain-containing protein, laterally wraps around and links the C1 and C2 microtubules. Furthermore, through single-molecule microtubule dynamics assays and cryo-electron tomography (cryo-ET) of SPEF1 binding to reconstituted microtubules, we demonstrated that SPEF1 recognizes and binds to the microtubule seam and contributes to microtubule stabilization. These structural and functional attributes of SPEF1 explain its critical role in ciliary assembly.

## Results

### High-resolution structure of the tip CP

To obtain a high-resolution structure of the tip CP, we isolated cilia from *Tetrahymena*, split the axoneme into separate microtubules, and prepared for single-particle cryo-EM. We developed a strategy to automatically pick substoichiometric tip CP particles (Fig. S1A, Materials & Methods) and obtained an 8-nm repeating structure with an overall resolution of 3.9 Å (Fig. 1B, Fig. S1A, B, Supplementary Movie 1). The map at this resolution confirmed the C1 and C2 microtubule seam assignment proposed previously on the basis of subtomogram averaging (Legal, Parra et al. 2023).

The C1 and C2 microtubules are almost symmetrical, with ∼180° rotation along the microtubule axis (Fig. 1B). The C1 and C2 microtubules are also linked by two asymmetric bridges (Fig. 1B). The major bridge spans between protofilaments (PFs) 3-4-5 of C1 and 1-13-12-11 of C2, whereas the minor bridge spans between PFs 12-13-1 of C1 and 3-4 of C2. In addition, there is a helical bundle on C2 close to the major bridge (Fig. 1B). We also identified two asymmetric densities on the microtubules: one on C1 and another on C2 (Fig. S1C). Three-dimensional classification of this helical bundle revealed a distinct 16 nm long feature (Fig. S1D), which contrasts with other densities that repeat every 8 nm.

Some of the symmetrical densities wedge between PFs, whereas others bind to the tubulin surface. These are the seam wedge density binding to the microtubule seam, Wedge density 1 positioned between PFs 11/12, Wedge density 2 between PFs 10/11 and Wedge density 3 between PFs 5/6. The tubulin surface density 1 is on PFs 8/9, and the tubulin surface density 2 is on PF 6. Inside the microtubule, there are eight filamentous microtubule-inner proteins (MIPs) (Fig. 1B, purple) and an MIP with an arc density spanning PF11-12-13-1 (Fig. 1B, yellow).

### Identification of proteins that bind specifically to the tip CP

While the density in the center region of the map was well resolved, the resolution outside the center region was insufficient to identify proteins *de novo* on the basis of side chain density. Therefore, we used DomainFit (Gao, Tong et al. 2024) and our database of axonemal proteins (Kubo, Black et al. 2023) to identify the domains associated with the densities binding to microtubules (Fig. S1E). By integrating the DomainFit results with *in situ* crosslinking-mass spectrometry of cilia (McCafferty, Papoulas et al. 2023) and the relative abundance of proteins in the axoneme, we were able to identify seven proteins in the tip CP (Fig. 1C, D and Supplementary Table 1). The remaining parts of the identified proteins were modelled via a combination of AlphaFold2-predicted models and manual modelling (Materials & Methods).

These analyses revealed that the seam wedge density is occupied by a CH domain from either SPEF1A or SPEF1B (TTHERM_00939230, TTHERM_00161270). Since we detected only SPEF1A in the proteome of the axoneme (Kubo, Black et al. 2023), we assigned only SPEF1A to the density. SPEF1B may also occupy the seam wedge density at a substoichiometric ratio compared with SPEF1A. The three other wedge densities are formed by two proteins containing tubulin polymerization-promoting protein (TPPP)-like domains, i.e., TPPP-like protein 1 (TLP1, TTHERM_00070880) and 2 (TLP2, TTHERM_00370820). TLP2 consists of two TPPP-like domains (Wedge density 2 and Wedge density 3) linked by a long helix spanning five PFs. The two tubulin surface densities are composed of a C2 domain-containing protein, namely, C2D1 (TTHERM_00243990) (Tubulin surface density 1), and a homodimer of protein with a small DPY-30 motif (TTHERM_00579000/TTHERM_00082260/TTHERM_00158390) (Tubulin surface density 2). The spike density was identified as a protein containing an LSDAT domain (TTHERM_000686039), which we named SPIKE1 (TTHERM_000686039). The arcMIP linking PF 11-1 is CFAP213 (TTHERM_00773430), a homologue of FAP213, which binds to similar PFs in the C2 microtubule in the main ciliary shaft of *Chlamydomonas* (Han, Rao et al. 2022), whereas the filamentous MIPs (Fig. 1C, purple) remain unidentified.

To verify that the correct proteins were identified, we expressed them as GFP fusions in *Tetrahymena* cells under the control of the respective native promoters. Using superresolution structural illumination microscopy (SR-SIM), we visualized the GFP-tagged proteins in conjunction with either the anti-monoglycylated antibody TAP952, which preferentially labels the ciliary tip and assembling cilia (Fig. 2A), or the antipolyglycylated antibody AXO49, which labels the main axoneme excluding the ciliary tip (Fig. S2).

**Figure 2:**
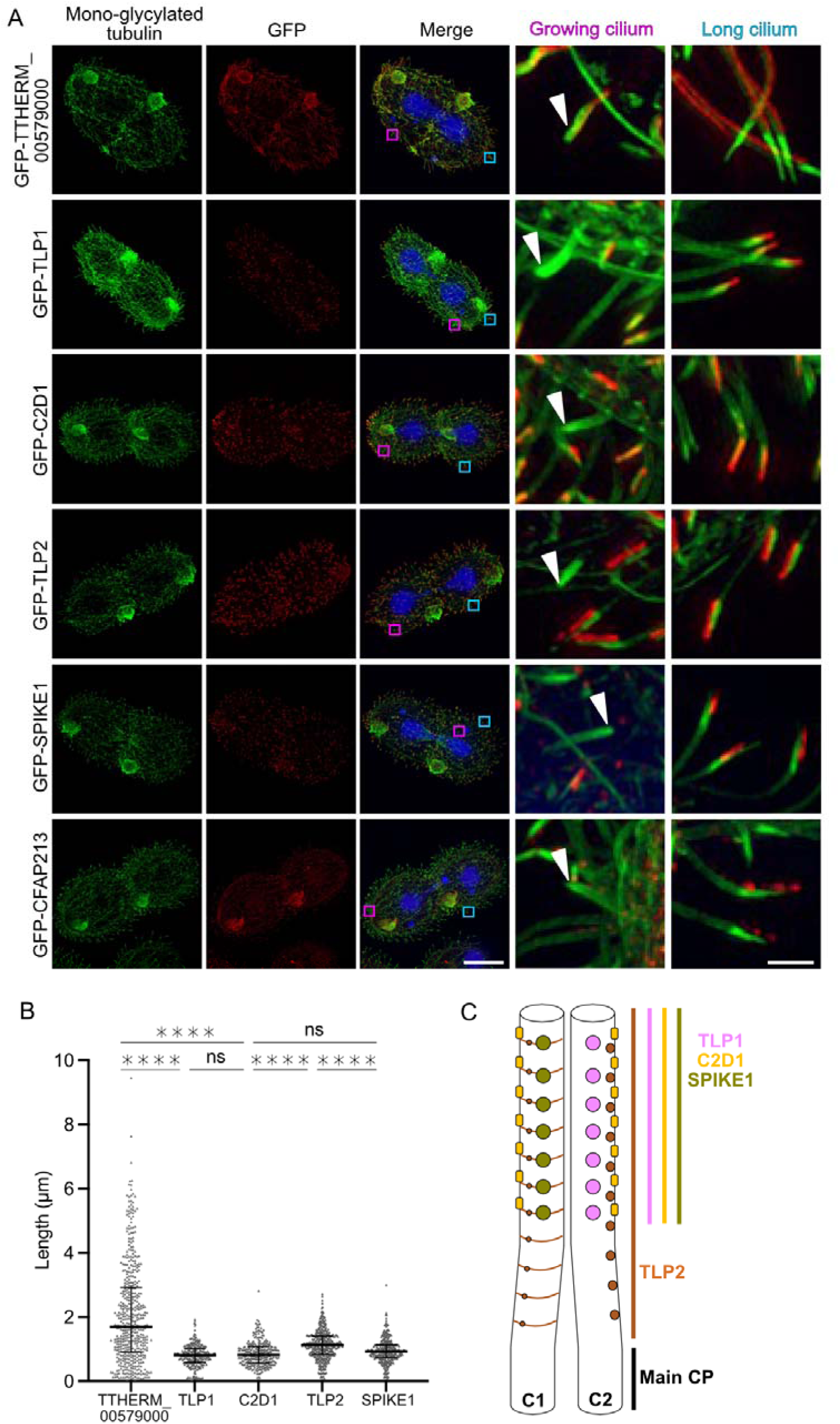
The proteins identified by DomainFit localize to the tips in vivo. (A) Representative fluorescence images of mono-glycylated tubulin (green), GFP-tagged TTHERM_00579000 (DPY-30 motif protein), TLP1, C2D1, TLP2, SPIKE1 and CFAP213 (red), and DNA (blue) and merged images. Scale bar: 20 μm. Insets show both a growing cilium (pink square in the merged image) and a long cilium (blue square in the merged image). Scale bar: 1.5 μm (B) Quantification of the signal length for each protein except for CFAP213. Number of measurements: n=485, 261, 278, 419 and 334 for DPY-30 (TTHERM_00579000), TLP1, C2D1, TLP2 and SPIKE 1, respectively. The horizontal bars indicate the medians. The error bars indicate the interquartile ranges. **** p<0.0001, Kruskal□Wallis test. CFAP213 fluorescence was not quantified because of its appearance as spots. (C) Diagram showing the distribution of the tip proteins along the length of the tip CP.

**Supplementary Figure 2:**
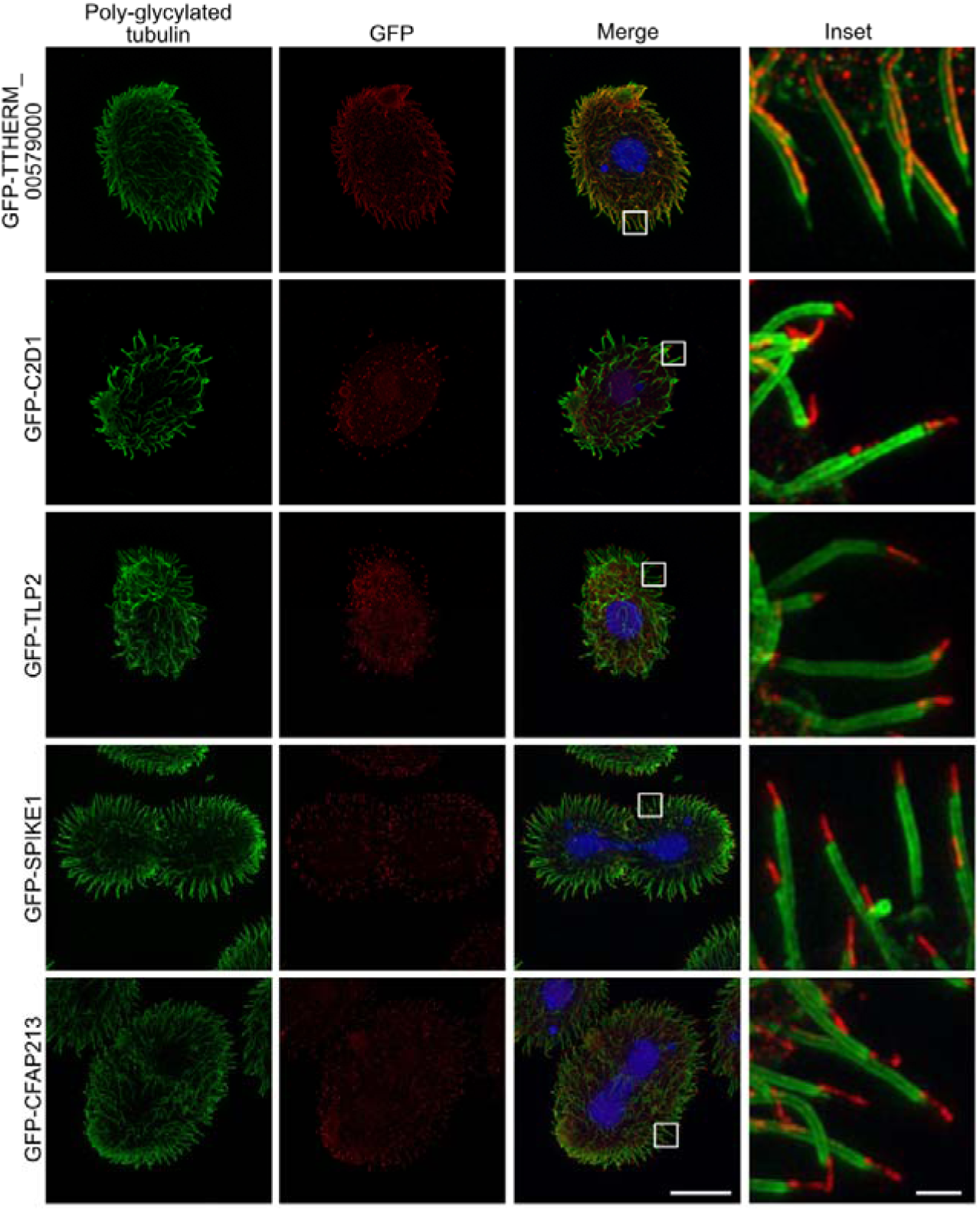
Representative images of the colocalization of polyglycylated tubulin (green) and GFP-tagged TTHERM_00579000, TLP1, C2D1, TLP2, SPIKE1 and CFAP213 (red) and DNA (blue). The last column shows merged images. Scale bar: 20 μm. Insets zoomed in on representative cilia from the cell shown in the merged image. Scale bar: 1.5 μm.

Except for the DPY-30 motif protein (TTHERM_00579000) (mapped to the Tubulin surface density 2), all the other proteins were located in the tip region. As the tagged TTHERM_00579000 is present along the main region of cilia, dimers of other DPY-30 motif-containing proteins (TTHERM_00082260 or TTHERM_00158390) may occupy the Tubulin Surface Density 2 instead (Fig. S1F). The signal from GFP-SPEF1A was undetectable, probably because the GFP tag interferes with SPEF1A binding to microtubules. On the other hand, SPEF1B was enriched in the tips of the cilia of mNeon-SPEF1B-expressing *Tetrahymena* cells (Guha, Vasudevan et al. 2024).

Labelling cilia with anti-monoglycylated antibodies in cells expressing GFP-tagged proteins allowed us to assess whether tip CP proteins were present in short, growing cilia in dividing cells (Fig. 2A). In growing cilia, we were unable to detect the GFP signal (localizing to the tip CP proteins) (except SPEF1A, for which data are not available) until the cilia were approximately half or two-thirds in length (Fig. 2A, white arrowheads).

The measurements of the length of the tip CP fragment occupied by the analysed proteins (the fluorescence GFP signal) revealed that SPIKE1, TLP1, and C2D1 span similarly ∼0.87 μm. In comparison, TLP2 occupies a longer region of ∼1.13 μm (Fig. 2B). This finding suggests that TLP2 binding starts in the transition region between the main and tip CP (Fig. 2C), whereas SPIKE1, TLP1, and C2D1 bind only in the tip region. The localization of TLP2 in a longer portion of the tip may suggest that TLP2 assembles first and then recruits other tip CP proteins.

### Characterization of TLP2 as a microtubule-associated protein spanning laterally across numerous PFs

TLP2 is a large protein of 1694 residues. It has three main globular domains: an N-terminal Armadillo repeat (ARM) domain (residues 143-706) and two TPPP-like domains (residues 889-1064 and 1489-1657) (Fig. 3A). A striking feature of TLP2 is that it spans laterally over eight PFs (Fig. 3B). Its TPPP-like domain 1 binds at the intradimer interface between PFs 5 and 6, whereas TPPP-like domain 2 binds between PFs 10 and 11, also at the intradimer interface. Both TPPP-like domains are linked by a long helix that contacts only β-tubulin on PFs 6 to 8 and binds at the intradimer interface of PFs 9 and 10 (Fig. 3B).

**Figure 3:**
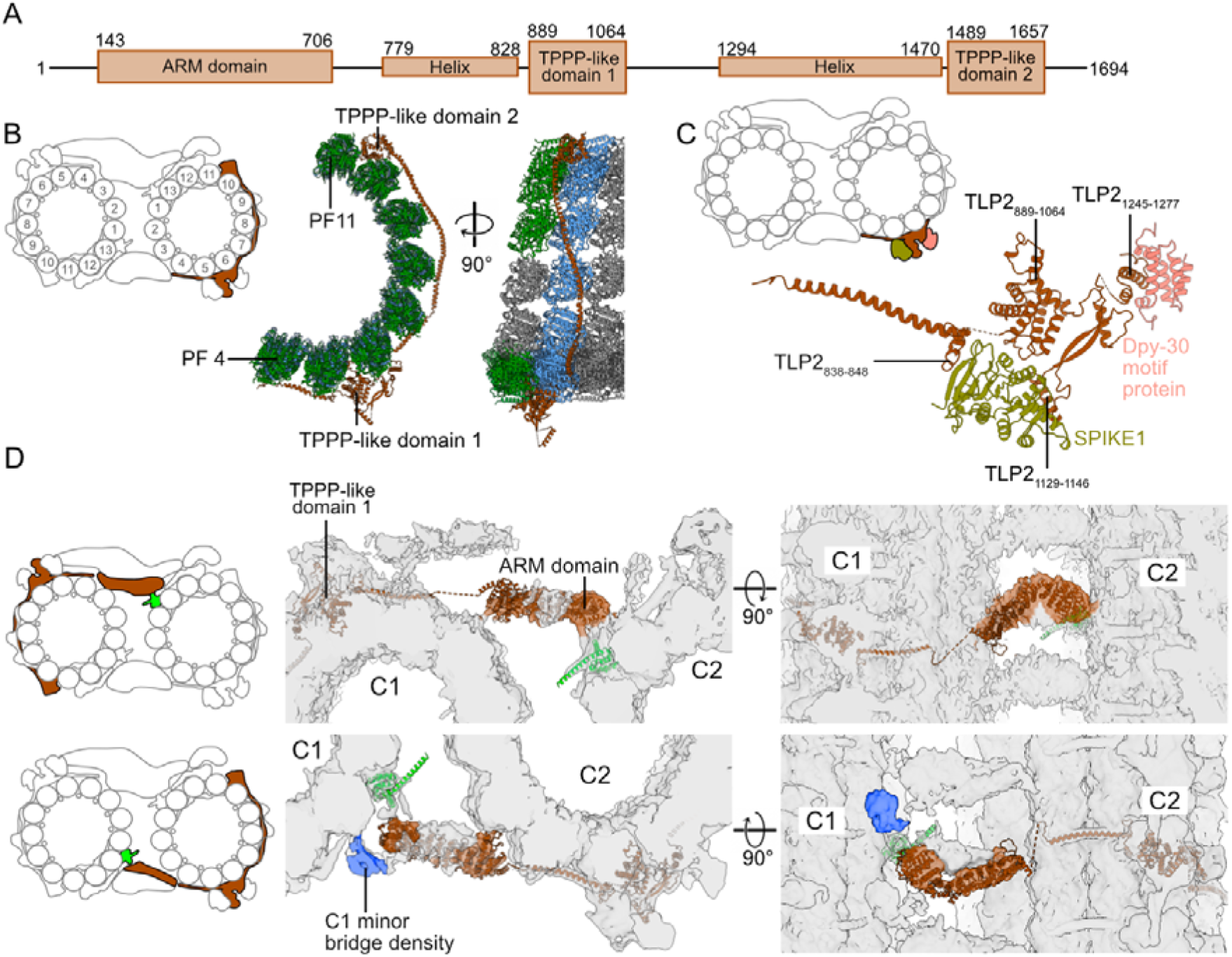
TLP2 binds eight PFs simultaneously. (A) Diagram showing the different domains of TLP2. (B) Atomic model of TLP2 bound to PFs 4 to 11 of C2. (C) Atomic model of the spike density. SPIKE1 binds to the TLP2 helices TLP2_838-848_ and TLP_1129-1146_ and the TLP2 TPPP-like domain TLP2_889-1064_. DPY-30 motif-containing protein binds to the TLP2 helix TLP2_1245-1277_. (D) Predicted model of the TLP2 ARM domain, TLP2_143-706_ fitted with the cryo-EM densities of both tip CP bridges. The density in blue is present only on C1 near the minor bridge.

**Supplementary Figure 3:**
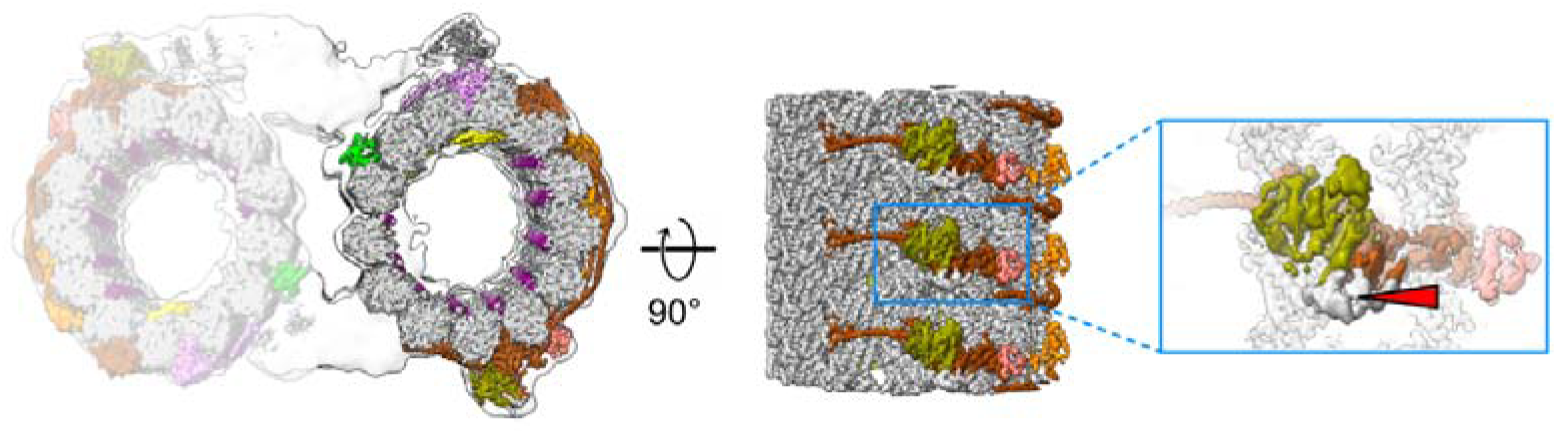
Spike density coloured near the predicted model of SPIKE1 (green) and TLP2 (brown). The red arrowhead points to a density whose protein was not identified (gray).

TLP2 acts as a scaffold for SPIKE1 and a dimer of DPY-30 motif proteins. These four proteins and another as-yet unidentified protein form the characteristic spike (Fig. 3C, Fig. S3). The protein SPIKE1 is positioned between TLP2 helix (residues 838-848) and TPPP-like domain 1. The DPY-30 motif protein binds to a helix of TLP2 formed by residues 1245-1277 (Fig. 3C).

We identified a low-resolution density of both major and minor bridges connecting C1 and C2 and matched the characteristic shape of the TLP2 ARM domain (Fig. 3D). Therefore, TLP2 likely contributes to the stability of the tip CP by binding to both C1 and C2. In addition, the ARM domain in the minor bridge is close to an unidentified C1 specific density (Fig. 3D). The ARM domains of TLP2 on both C1 and C2 microtubules are located close to SPEF1 and thus can bind to SPEF1 either directly or via an accessory protein.

While there are a few known α-helical proteins that bind to microtubules, such as MAP7 (Ferro, Fang et al. 2022), and many filamentous MIPs, such as CFAP45, CFAP112, CFAP53 and CFAP210 (Ma, Stoyanova et al. 2019, Kubo, Black et al. 2023), TLP2 is the first α-helical protein reported to extend laterally across the microtubule surface. With these expansive contacts with the microtubule surface by the helix connecting the two TPPP-like domains, TLP2 can strongly stabilize C1 and C2 microtubules at the ciliary tip.

### Preference of TLP1, TLP2, and CFAP213 for high inter-PF curvature

TLP1, another tip protein, also contains a TPPP-like domain (Fig. 4A). In mammals, three paralogues of TPPP are expressed in different tissues. They all have a globular domain whose predicted structures are similar, followed by a flexible region, with TPPP1 also possessing an additional N-terminal flexible extension. TPPP2 is involved in spermiogenesis and results in decreased sperm motility in KO mice (Zhu, Yan et al. 2019).

**Figure 4:**
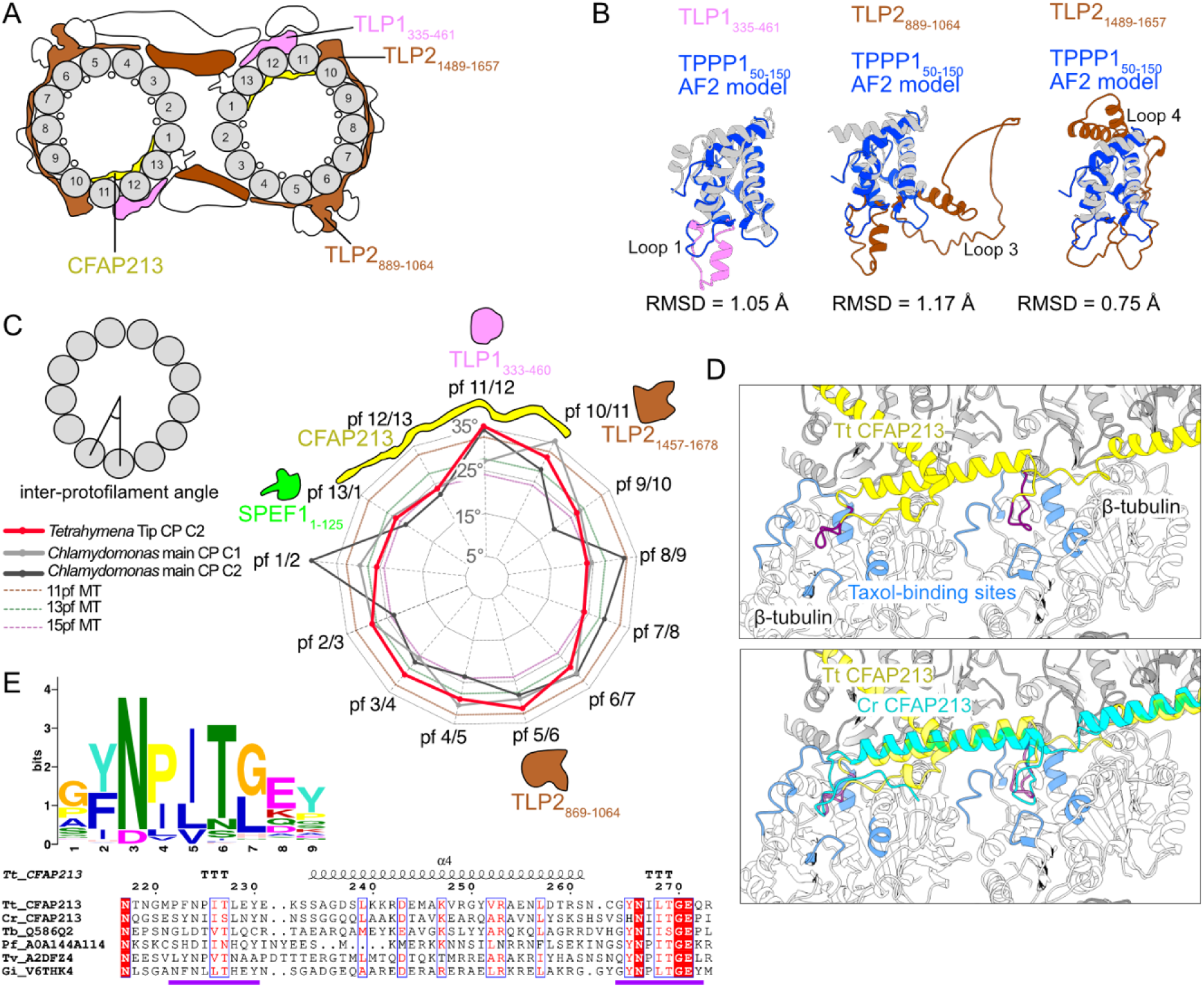
TPPP-like domains bind to high-curvature regions of microtubules. (A) Diagram of the tip CP highlighting the TLP1 and TLP2 TPPP-like domains and CFAP213. (B) Overlay of the HsTPPP1_50-150_ and TPPP-like domains of TLP1 and TLP2. (C) Inter-PF angles of Tetrahymena tip CP C2 microtubules, Chlamydomonas main CP microtubules and 11-, 13- and 15-PF microtubules. (D) Overlay of Tetrahymena (yellow) and Chlamydomonas (cyan) CFAP213, showing the loop (purple) that interacts with the taxane-binding sites (blue) on β-tubulin. (E) Logos of multiple sequence alignments of the predicted loops in MIPs. Parts of the alignment of Tetrahymena (Tt), Chlamydomonas (Cr, UniProtID A8JB78), Trypanosoma bruceii (Tb, UniProtID Q586Q2), Plasmodium falciparum (Pf, UniProtID A0A144A114), Trichomonas vaginalis (Tv, UniProtID A2DFZ4) and Giardia intestinalis (Gi, UniProtID V6THK4) CFAP213 homologues. The purple line marks the taxane pocket-interacting loops.

**Figure S4:**
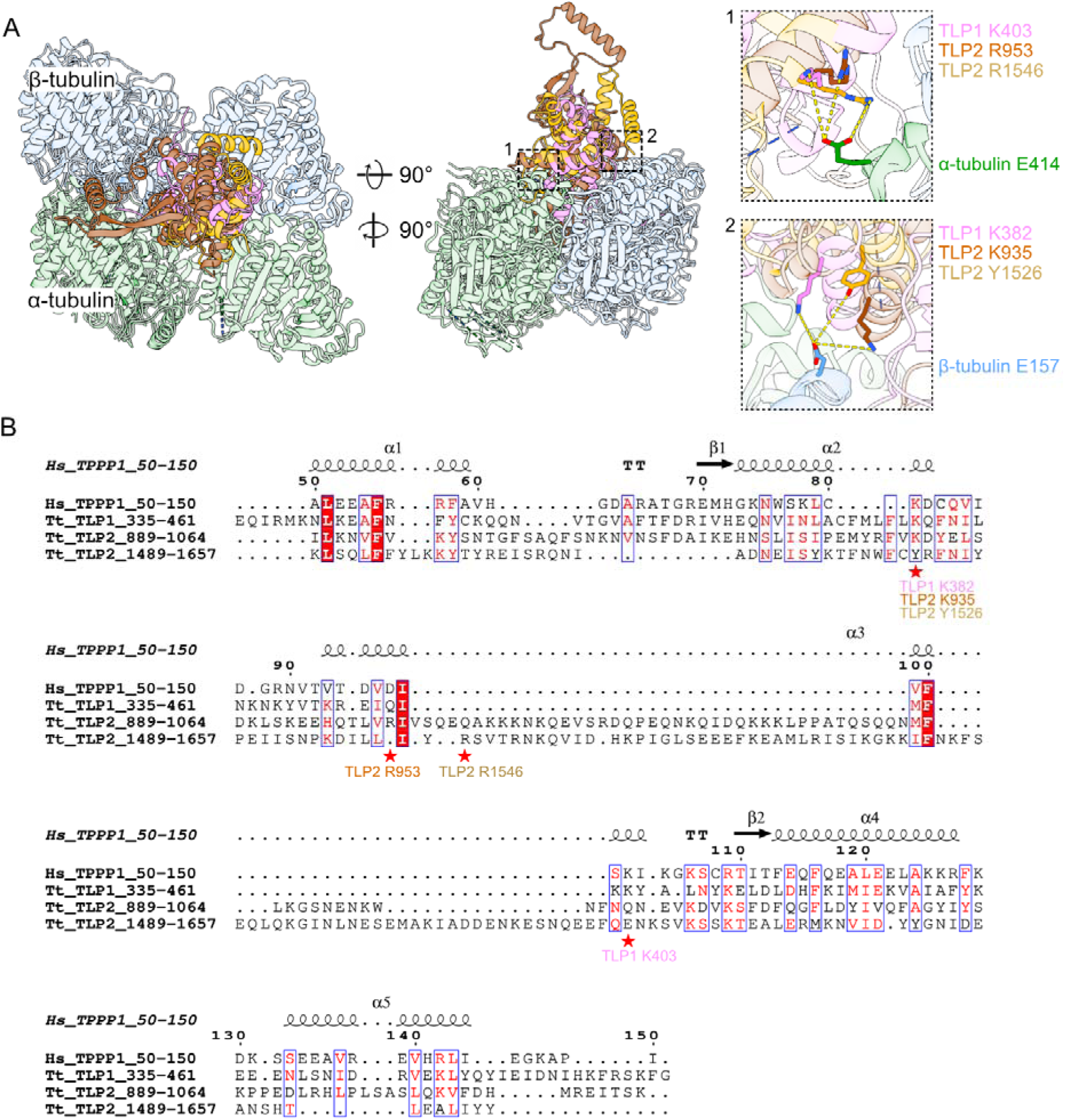
(A) Models of TPPP-like domains of TLP1 and TLP2 interacting with tubulin. (B) Alignment of the human TPPP1 (UniProtID O94811) and TPPP-like domains of TLP1 and TLP2. The amino acids shown in panel A are indicated by a red star.

The TPPP-like domains of TLP1 and TLP2 overlay most helices of the AlphaFold-predicted structure of the globular domain of human TPPP1_50-150_ but have longer loops and some extra helices. The TLP1_335-461_ fragment has a longer loop 1 with a small α-helix. The TLP2_889-1064_ fragment has a loop 1 similar to that of TLP1_335-461_ but has a much longer loop 3 made of residues 956-1019. Compared with human TPPP1, the TLP2_1489-1657_ region has slightly longer loops 1 and 3 and has a long extension around loop 4 made by residues 1579-1647 (Fig. 4B).

We measured the inter-PF angles of the tip CP microtubules and found that TPPP-like domains preferentially bind between PFs with high inter-PF angles (Fig. 4C). Compared with C1 and C2 from the main CP in *Chlamydomonas*, both C2 in the main and tip CP have remarkably high inter-PF angles between PFs 11 and 12 (34.5° and 35.3°, respectively; Fig. 4C). By overlaying the structures between two tubulin dimers and looking at their sequence alignment (Fig. S4A, B), we identified two sites with conserved residues among the TPPP-like domains that likely mediate the interaction with tubulins (Fig. S4A). TLP1 K403 and TLP2 R953 and R1546 are conserved and likely interact with α-tubulin E414, whereas TLP1 K382 and TLP2 K935 and Y1526 likely interact with β-tubulin E157. It is unclear whether the TPPP-like domains prefer to bind to high microtubule curvature or induce the formation of high curvature after binding.

In the high-curvature region between PFs 10 and 13, CFAP213 binds inside the microtubule lumen. Its homologue, CrFAP213, in *Chlamydomonas* is found only in the C2 microtubule of the main CP (Han, Rao et al. 2022). Since the tip CP microtubules have a similar curvature distribution to the C2 microtubule of the main CP, CFAP213 is likely the inducer of the unusual curvature distribution in the CP (Fig. 4C).

CFAP213 contains two loops that bind to the taxane-binding pockets of β-tubulin from PFs 12 and 13 (Fig. 4D). We found that these two loops are highly conserved. Using an HMM search with profiles built from TtCFAP213 and CrFAP213, we identified homologues of CFAP213, mainly from protists, including many parasites. We did not find any homologues of such proteins in higher eukaryotes, which raises a very interesting question as to whether alternative MIP motifs are used in these organisms to promote highly curved PFs. Using the motif discovery tool MEME (Bailey, Johnson et al. 2015), we identified the motif covering both loops that bind to the taxane-binding pocket as GYNPITGEY (Fig. 4E). CFAP213 is likely essential for the integrity of the tip CP, as observed in another MIP that also binds to the taxane-binding pocket Rib43a (Ichikawa, Khalifa et al. 2019).

### Characterization of SPEF1 binding to the microtubule seam

SPEF1 binds the microtubule seam via its CH domain (Fig. 5A). SPEF1A is more abundant in cilia than SPEF1B in mass spectrometry of cilia (Kubo, Black et al. 2023). Therefore, SPEF1A likely represents the majority of the seam-bound SPEF1 densities in the tip CP. Consistent with this observation, the density corresponding to SPEF1 includes the unique longer portion of helix 6, which is not present in SPEF1B, and other CH domain-containing proteins in the *Tetrahymena* axoneme (Fig. S5A). In Chlamydomonas, a seam-bining protein, CrFAP178, is found only on the seam of the C2 microtubule in the main CP and shares an almost identical CH-fold with SPEF1A (RMSD = 1.23 Å) (Gui, Wang et al. 2022, Han, Rao et al. 2022) (Fig. S5B, C).

**Figure 5:**
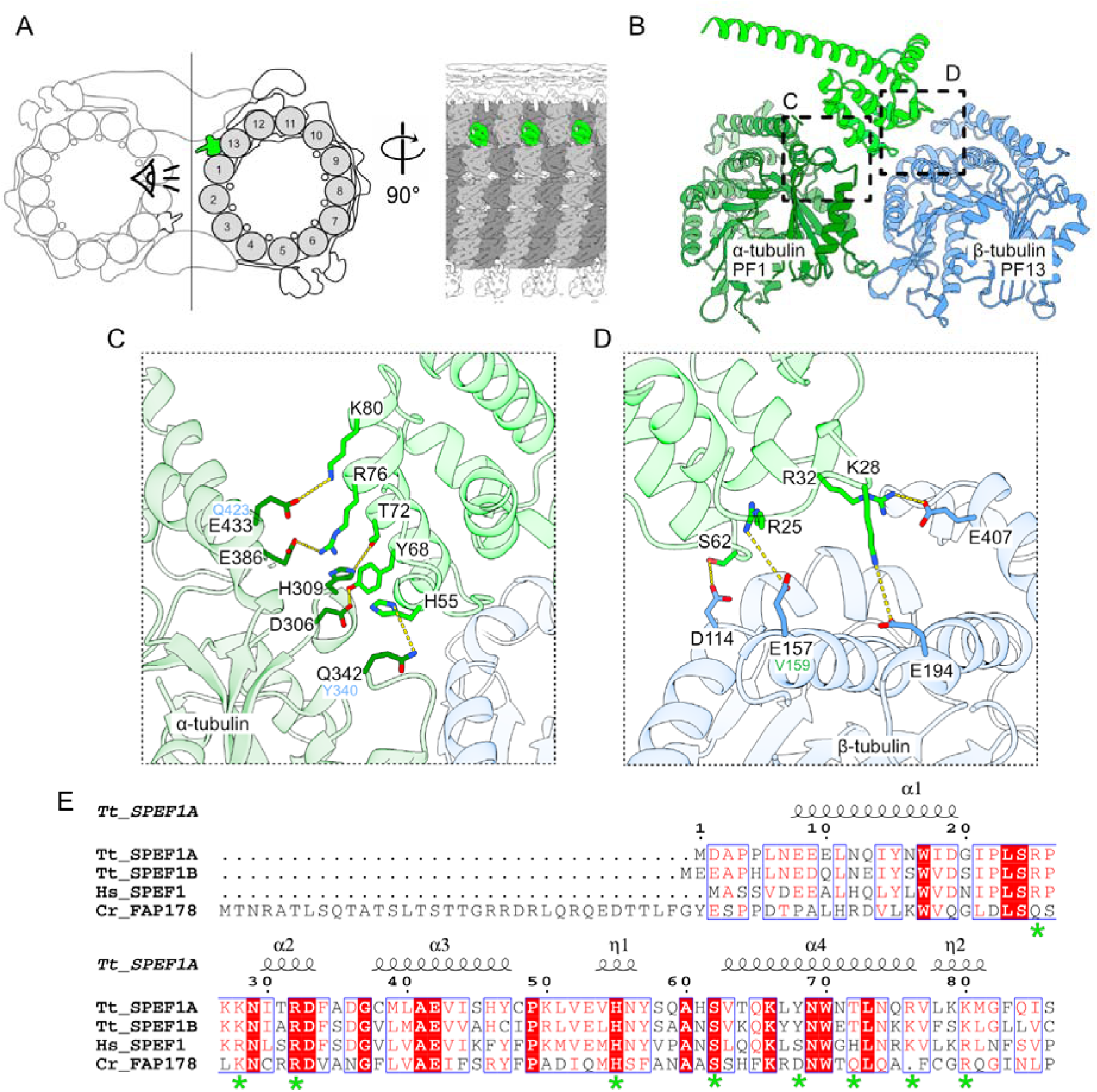
SPEF1 is a seam-binding protein. (A) Diagram of the tip CP highlighting SPEF1A (green). α-Tubulin is light gray, and β-tubulin is dark gray. (B) Model of SPEF1A interacting with tubulin at the seam. (C, D) Amino acids that likely mediate the interaction between α-(C) and β-(D) tubulin. The equivalent amino acids in β-and α-tubulin are shown in blue in (C) and in green in (D), respectively. (E) Alignment of Tetrahymena SPEF1A_1-86_, SPEF1B, human SPEF1 (UniProtID Q9Y4P9) and Chlamydomonas FAP178 (UniProtID A0A2K3D8Z6). The amino acids shown in C and D are highlighted with a green asterisk.

**Figure S5:**
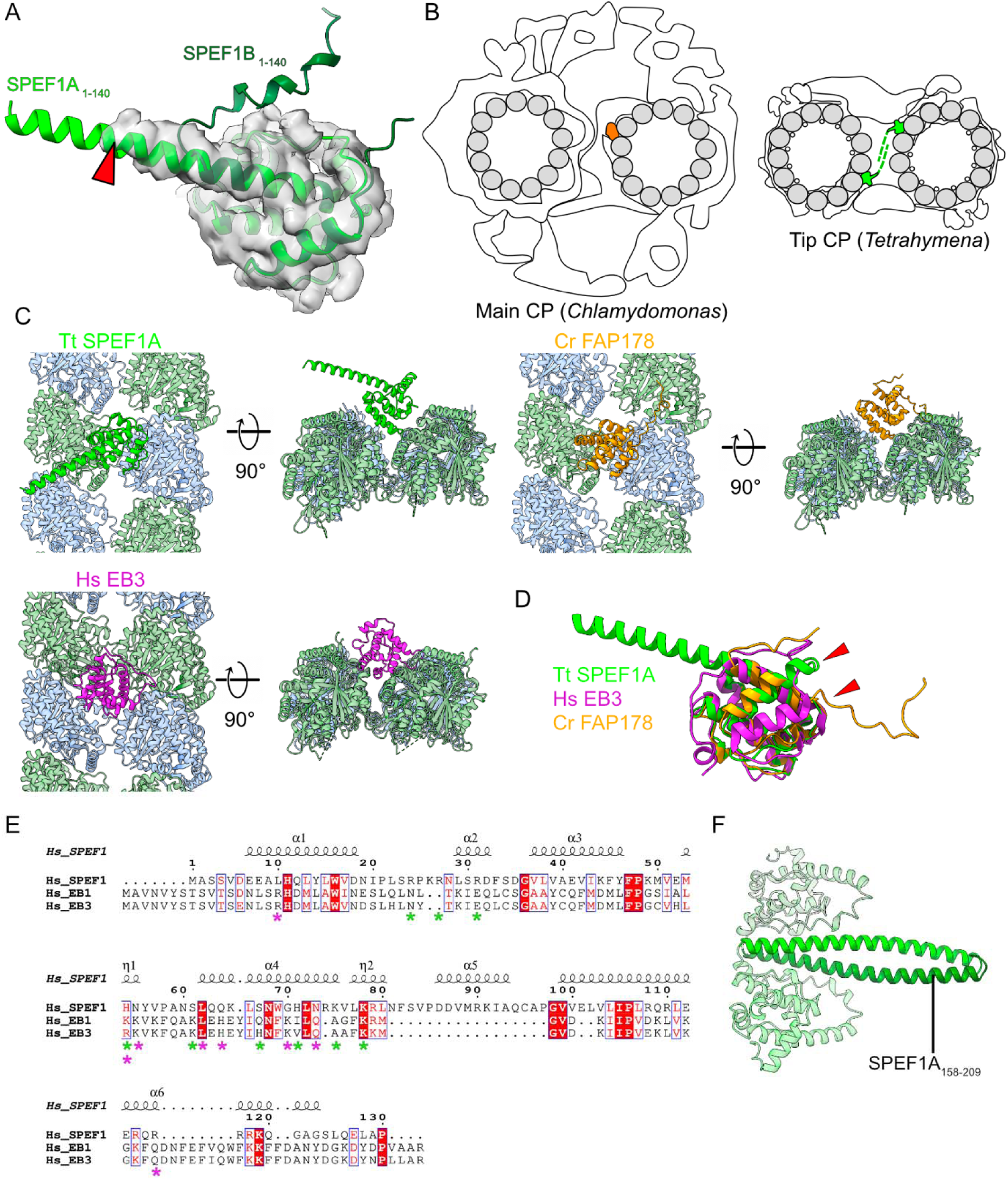
(A) Overlay of SPEF1A_1-140_ and SPEF1B_1-140_. The SPEF1 density from the CP map is in gray. The red arrowhead highlights helix 6 of SPEF1A. (B) Diagrams of the main (left) and tip (right) CPs showing that SPEF1A might dimerize between C1 and C2. CrFAP178 is shown in orange. SPEF1A is shown in green. (C) Models of TtSPEF1A (green), CrFAP178 (orange, EMDB-25361) and HsEB3 (pink, EMDB-6365) interacting with tubulin. (D) Overlay of TtSPEF1A, HsEB3 and CrFAP178. The arrowheads point to a different orientation of helix 1. (E) Alignment of human HsSPEF1_1-130_ (UniProtID Q9Y4P9), EB3 (UniProtID Q9UPY8) and EB1 (UniProtID Q15691). The amino acids interacting with tubulin are indicated by a purple asterisk for EB3 and a green asterisk for SPEF1A. (F) Predicted Alphafold2 model of a SPEF1A dimer. The helix mediating dimerization (SPEF1A_158--209_) is highlighted in green.

In *Tetrahymena*, SPEF1A binds α-tubulin of PF 1 and β-tubulin of PF 13 (Fig. 5B). Our model suggests that SPEF1A helix 4 interacts with α-tubulin. Positively charged residues R76 and K80 of SPEF1A likely make electrostatic interactions with E386 and E433 on α-tubulin helices 11 and 12, respectively. Moreover, Y68 and T72 on SPEF1A helix 4 likely interact with D306 and H309 positioned between helix 9 and beta strand 8 on α-tubulin. Finally, SPEF1A H55 likely interacts with Q342 on α-tubulin (Fig. 5C).

The interaction between SPEF1A and β-tubulin appears to be electrostatic and involves four key residues, R25, K28, R32, and S62, on SPEF1A, which likely interact with E157, E194, E407, and D114, which are located on helices 4, 5, 11, and 3 of β-tubulin, respectively (Fig. 5D). The residues mediating the interaction are conserved across species, suggesting a conserved mechanism of interaction (Fig. 5E). We believe that seam recognition is mediated by residues E433 and Q342 in α-tubulin, as these residues correspond to Q423 and Y340 in β-tubulin as well as E157 in β-tubulin, which corresponds to V159 in α-tubulin (Fig. 5C, D). Consistent with our findings, a previous study showed that in mouse SPEF1, mutation of the R31 residue (equivalent to R32 in *Tetrahymena*) to alanine impaired the ability of SPEF1 to bind and bundle microtubules when it was overexpressed in U2OS cells and could not rescue CP formation or ciliary motility in SPEF1-depleted mouse ependymal cells (Zheng, Liu et al. 2019).

CH domains are also present in end-binding proteins (EB1 and EB3) to bind microtubules at the interdimer interface of four adjacent tubulin dimers except at the seam (Fig. S5C) (Maurer, Fourniol et al. 2012, Zhang, Alushin et al. 2015). Thus, there is a stark contrast in how the CH domains present in SPEF1 and EBs bind to microtubules. While the folds of the CH domains in SPEF1A and EB3 are similar, helix 1 has a markedly different conformation (Fig. S5D, red arrowheads). In addition, SPEF1A-microtubule binding involves amino acid residues that are not conserved in EB3 or EB1 and vice versa (Fig. S5E).

In mice, SPEF1 was previously shown to form a homodimer through its C-terminal helix (Zheng, Liu et al. 2019). AlphaFold2 prediction of a *Tetrahymena* SPEF1A dimer indicated a confident prediction, with a pTM of 0.83 for SPEF1A_158-209_ corresponding to helix 7 (Fig. S5F). However, we did not observe any density corresponding to the dimerization domain of SPEF1A. Since the distance between two *Tetrahymena* SPEF1A molecules facing each other on C1 and C2 is ∼8 nm, similar to the repeating length of SPEF1A on each microtubule, SPEF1A can therefore dimerize on the same microtubule or crosslink C1 and C2 microtubules (Fig. S5B). This arrangement is different from that of CrFAP178, which solely exists on C2 microtubules and thus cannot crosslink C1 and C2 microtubules.

### Binding of the human SPEF1 CH domain to the seam and stabilization of microtubules *in vitro*

The microtubule seam is believed to be a weak point in the microtubule lattice, as lateral contacts are formed between α- and β-tubulin (Alushin, Lander et al. 2014). As *Tetrahymena* SPEF1A recognizes this unique interaction in the seam, we hypothesized that human SPEF1 could function similarly and stabilize microtubules. We purified a fragment of human SPEF1 containing a CH domain fused to a GFP tag (HsSPEF1-CH) (Fig. S6A) and used cryo-ET to visualize Taxol-stabilized reconstituted microtubules incubated with HsSPEF1-CH. Notably, we observed many microtubule pairs, similar to the native tip CP, suggesting that HsSPEF1-CH can crosslink both microtubules despite lacking the C-terminal dimerization domain (Fig. 6A). We also observed microtubule bundles (Fig. S6B). These findings suggest that HsSPEF1-CH can dimerize, possibly through its GFP tag (Phillips 1997).

**Figure 6:**
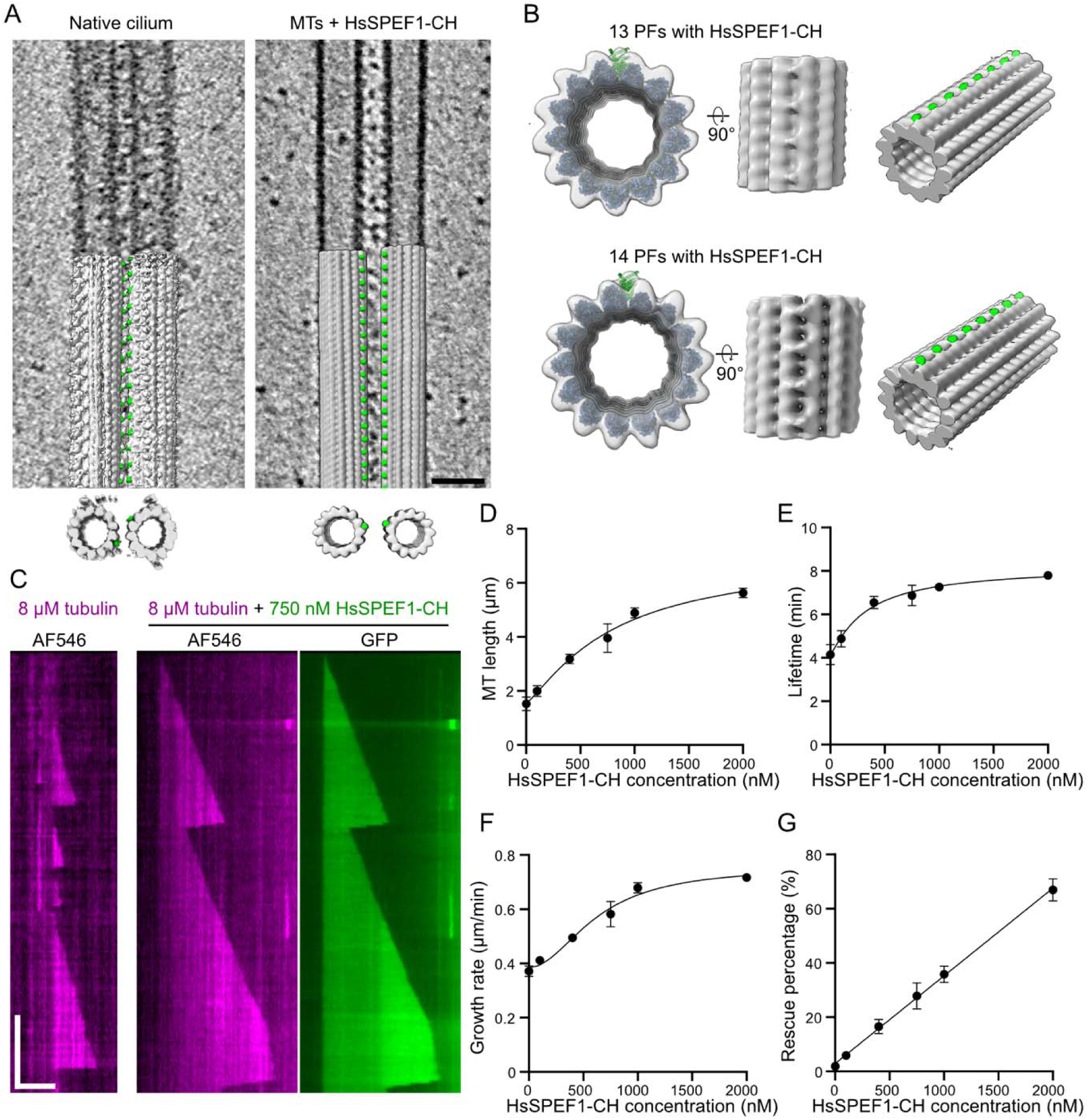
SPEF1 stabilizes microtubules by binding to the seam. (A) Denoised (SIRT-like filter) tomogram of purified tip CP (native cilium data from Legal *et al*., 2023) and microtubules incubated with HsSPEF1-CH. Arrowheads point to a repetitive density found between both microtubules. Scale bar: 40 nm. (B) Subtomogram averaging map of the 13-PF and 14-PF microtubules incubated with HsSPEF1-CH overlapping with models of 13-PF and 14-PF microtubules binding to HsSPEF1-CH. The density corresponding to HsSPEF1-CH is coloured in green. (C) Representative kymographs of 750 nM recombinant HsSPEF1-CH and 8 μM MT. Horizontal scale bar: 4 μm; vertical scale bar: 5 minutes. (D - G) Microtubule length, lifetime, growth rate and rescue probability versus HsSPEF1-CH concentration. Each point represents the average, and the error bars represent the standard deviation. n=421, 482, 568, 643, 580, and 555 microtubules for HsSPEF1-CH concentrations of 0, 100, 400, 750, 1000, and 2000, respectively, obtained from three independent experiments.

**Figure S6:**
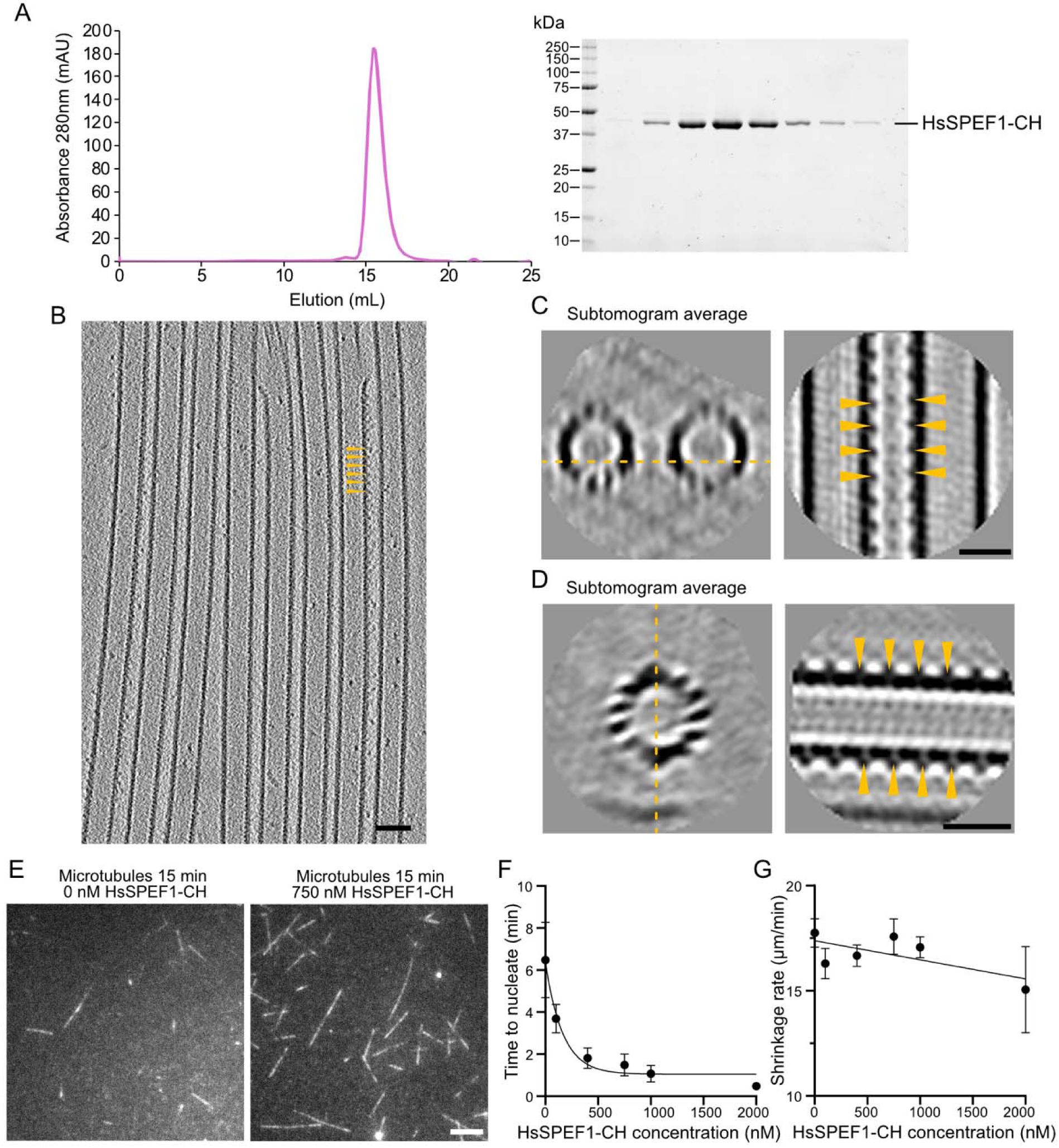
(A) Curve after SEC and the corresponding gel for HsSPEF1-CH. (B) Example tomogram of microtubules with HsSPEF1-CH. Arrowheads point to extradensity binding to the side of a microtubule with an apparent repeat of 8 nm. Scale bar, 40 nm. (C - D) Subtomogram averages of a microtubule pair shown in Fig. 6F and a single microtubule from another tomogram. Arrowheads point to extra densities that repeat every 8 nm. Scale bar 20 nm. (E) Representative TIRF images of the AF546 channel (microtubules) at t = 15 min with and without 750 nM HsSPEF1-CH. Scale bar, 4 μm. (F) Time for microtubules to nucleate vs the HsSPEF1-CH concentration. (G) Microtubule shrinkage rate vs the HsSPEF1-CH concentration.

Subtomogram averaging revealed a mix of 13- and 14-PF microtubules. Both showed an 8-nm repeat density only between one PF pair, which has not been observed in any other microtubule-associated proteins (Fig. 6B, S6C and Supplemental Movie 2). While the resolution of the subtomogram average is not high enough to visualize the seam, this unique binding pattern strongly suggests that HsSPEF1-CH bind to the seam *in vitro* (Fig. 6B). Occasionally, we observed extra density along different PFs on the same microtubule (Fig. S6D). As microtubules are Taxol stabilized, we suspect that those microtubules contain multiple seams that are recognized by HsSPEF1-CH (Kikkawa, Ishikawa et al. 1994, Debs, Cha et al. 2020, Guyomar, Bousquet et al. 2022).

We then tested the ability of HsSPEF1-CH to stabilize growing microtubules *in vitro* via single-molecule assays as a seam binder (Fig. 6C, S6E). HsSPEF1-CH bound uniformly on dynamic microtubules and GMPCPP-stabilized seeds, suggesting that HsSPEF1-CH binds microtubules regardless of the nucleotide state. The binding of HsSPEF1-CH increased the microtubule length, lifetime, growth rate and rescue percentage in a concentration-dependent manner (Fig. 6D-G). HsSPEF1-CH also decreased the microtubule nucleation time but did not seem to affect the shrinkage rate (Fig. S6F-G). These data show that the HsSPEF1 CH-domain alone is sufficient to stabilize microtubules.

In summary, these data suggest that SPEF1 stabilizes and crosslinks the microtubules of the tip CP and might also regulate the distance between both CP microtubules.

## Discussion

We present the high-resolution structure of the CP microtubules in the tip region and identified seven new tip proteins, including SPEF1, CFAP213 and proteins containing TPPP-like domains. Furthermore, our *in vitro* experiments indicate that SPEF1 is a seam-binding protein that stabilizes and crosslinks tip CP microtubules.

Our structure suggests that the tip CP is highly stabilized by CFAP213 and TLP2. Our structure demonstrates how the TPPP-like domain interacts with high-curvature microtubules. The TPPP protein is found in the cilia of many species (Orosz 2024), but the role of the TPPP protein in cilia is unclear. In *Chlamydomonas*, knockout of TPPP does not affect ciliary motility but affects flagellar reassembly and results in hatching defects (Tammana and Tammana 2017). Therefore, we suspect that TLP1 and TLP2 might be important for cilia reassembly.

Our data show that SPEF1 binds to the seam *in vivo* and *in vitro* and potentially dimerizes to form parallel microtubules. Prior structural studies suggested that the microtubule seam is its weakest region, likely because the heterotypic lateral contacts across the seam are affected by dimer repeat changes associated with E-site remodelling during GTP hydrolysis (Alushin, Lander et al. 2014). Microtubule stabilization could be achieved through regularization of the interaction between PFs on either side of the seam, either by a microtubule-binding protein such as EB3 (Zhang, Alushin et al. 2015) or a drug that binds between PFs (Kellogg, Hejab et al. 2017). Here, using cryo-ET, we showed that SPEF1 binds specifically to the seam and stabilizes microtubules, as revealed by a single-molecule assay. As a result, our work confirms that seam stabilization improves microtubule stability. Unfortunately, owing to low resolution, we were unable to confirm any seam regularization in our cryo-ET data.

In addition to cilia, SPEF1 also localizes in other places, such as in pillar cells of the mouse cochlea, which maintain large extracellular spaces between them and contain extensive networks of highly ordered microtubule bundles (Dougherty, Adler et al. 2005). The ability of SPEF1 to recognize the seam and form a dimer likely enables the formation of bundles with uniformly oriented microtubules, as in the case of the CP.

The striking difference in composition and morphology between the tip and main CP raises questions regarding CP assembly and the role of SPEF1 in this process. Previous studies indicate that the CP assembly process is rather complex. Knockout of SPEF1 leads to improper assembly of the main CP in mouse ependymal cells and *Tetrahymena* (Zheng, Liu et al. 2019, Guha, Vasudevan et al. 2024). Although SPEF1 is localized only to the tip region of *Tetrahymena* and *Xenopus* (Konjikusic, Lee et al. 2023, Guha, Vasudevan et al. 2024, Rao, Subramanianbalachandar et al. 2024), it is transported to cilia immediately after deciliation and then gradually accumulates in the tip region in *Xenopus* (Rao, Subramanianbalachandar et al. 2024). Therefore, SPEF1 likely participates in the early stages of CP assembly. We propose that SPEF1 dimerization and seam binding ability can affect CP assembly either through proper tip organization or stabilization of the CP microtubules during assembly (Fig. 7).

**Figure 7:**
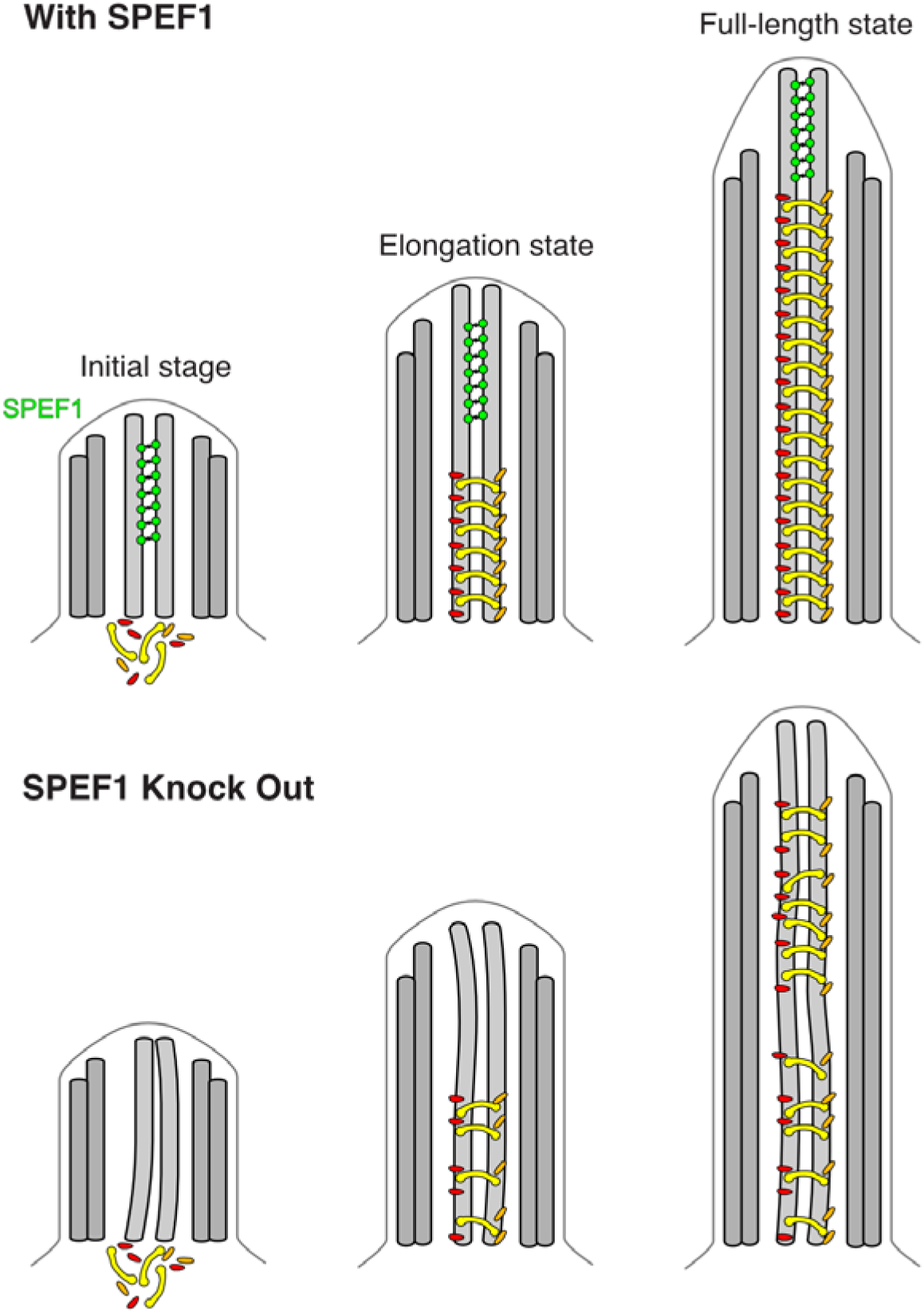
Model of the role of SPEF1 in CP stability. SPEF1 stabilizes the tip CP MTs from the start of their assembly, allowing the main CP to be correctly assembled. If SPEF1 is knocked out, the CP MTs are destabilized, and the main CP cannot assemble correctly, leading to defects in cilia function.

The functions of other nonconserved ciliary tip proteins, such as the C2D1 and SPIKE1 proteins, are unknown. C2 domain-containing proteins are generally found in cilia and centrioles. Interestingly, mutations in the C2 domain-containing and centriole protein CC2D2A cause Joubert syndrome (Gorden, Arts et al. 2008). As C2 domains are known to bind membranes, the C2 domain found in the tip may be used to bridge microtubules and the membrane and constrict the space in the tip. The constriction of space in the tip might be important for the remodelling of IFT to convert from anterograde to retrograde transport (Nievergelt, Zykov et al. 2022).

The structural diversity of the ciliary tip across species and tissues raises questions about its specific role. Our findings highlight a potential conserved function of SPEF1, suggesting a unifying theme in tip-mediated cilia assembly despite structural variations. However, further research is needed to elucidate the roles of tip proteins in this process.

## Materials and methods

### Tetrahymena culture

All *Tetrahymena* strains used in this study were grown in SPP media (Williams et al., 1980) in a shaker incubator at 30°C and 120 rpm. Cultures were harvested at an OD_600_ of approximately 0.6.

### Cilia isolation and preparation for cryo-EM

Cilia from the *Tetrahymena* CU428 strain were isolated as previously described (Kubo, Black et al. 2023). The cell cultures were harvested via centrifugation at 2000□×□*g* for 10□min at 23°C. Pelleted cells were resuspended in fresh SPP and adjusted to a total volume of 24□mL. Dibucaine (25□mg in 1□mL SPP) was added, and the culture was gently swirled for 45□s. To stop the dibucaine treatment, 75□mL of ice-cold SPP supplemented with 1□mM EGTA was added, and the dibucaine-treated cultures were centrifuged for 15□min at 2000□×□*g* and 4°C. The supernatant (cilia) was collected and centrifuged at 30,000□×□*g* for 45□min at 4°C. The pelleted cilia were gently washed a few times and resuspended in Cilia Wash Buffer (50□mM HEPES at pH 7.4, 3□mM MgSO4, 0.1□mM EGTA, 1□mM DTT, 250□mM sucrose). The resuspended cilia were flash-frozen with liquid nitrogen and stored at −80°C.

The frozen cilia were thawed on ice and then centrifuged for 10□min at 8000□×□*g* at 4°C. The pellet (cilia) was washed and resuspended in Cilia Final Buffer (50□mM HEPES at pH 7.4, 3□mM MgSO4, 0.1□mM EGTA, 1□mM DTT, 0.5% trehalose). Next, the NP-40 alternative (Millipore Sigma #492016) was added to a final concentration of 1.5%, and the cilia were incubated for 45□min on ice to remove the ciliary membrane. The de-membranated axonemes were centrifuged (8,000 × *g*, 4°C). The intact axoneme pellet was resuspended in Cilia Final Buffer with ADP (a final concentration of 0.3□mM). The sample was incubated at room temperature for 10□min. ATP was subsequently added to a final concentration of 1□mM, and the sample was again incubated at room temperature for 10□min. Samples were adjusted to 2.2□mg/mL using Cilia Final Buffer.

### Cryo-EM of the tip CP

C-Flat Holey thick carbon grids (Electron Microscopy Services #CFT312-100) were negatively glow-discharged (10□mA, 10□s). Four microliters of the cilia suspension were applied to the grids inside the Vitrobot Mk IV (Thermo Fisher) chamber. The sample was incubated on the grid for 15□s at 23□°C and 100% humidity, blotted with force 3 for 5□s, and then plunged frozen in liquid ethane.

### Cryo-EM data acquisition

A total of 18,384 movies (dataset from (Kubo, Black et al. 2023)) and 20,050 newly recorded movies were collected for the *WT* in two separate datasets using a Titan Krios 300□keV FEG electron microscope (Thermo Fisher Scientific) equipped with a direct electron detector K3 Summit (Gatan, Inc.) and a BioQuantum energy filter (Gatan, Inc.) using SerialEM. The movies were collected with a beam shift pattern of four movies per hole and four holes per movement. The final pixel size is 1.370□Å/pixel. Each movie had a total dose of 45 electrons per Å^2^ over 40 frames. The defocus range was between −1.0 and −3.0 μm, with an interval of 0.25 μm.

### Data processing

Data processing was performed using CryoSPARC v4.5.1 (Fig. S1A) (Punjani, Rubinstein et al. 2017). Patch motion correction and patch CTF were first applied to the movies. Particles in ∼1,000 micrographs were picked manually using e2helixboxer (Tang, Peng et al. 2007). Particles were extracted every 8 nm and used to train Topaz (Bepler, Morin et al. 2019) (box size 680 pixels, binned to 170 pixels). The remaining particles were picked via Topaz. The micrographs were manually curated to contain between 10 and 2000 particles, a CTF fit between 0 and 8 μm, and a relative ice thickness between 0 and 1.1. The particles were then subjected to two rounds of 2D classification. Helical refinement was then carried out using initial reference from subtomogram averaging of the tip CP (Legal, Parra et al. 2023). Interestingly, C2 had better resolution than C1. Local refinement was carried out on the central section of the map (approximately 24 nm). The particles were shifted +82 and −82 nm, and their center was changed to 0 before removing duplicates and bad particles through 2D classification. Particles were extracted using a box size of 680 pixels binned to 540. Local CTF refinement and reference-based motion correction were then carried out. Masks of 3 to 4 PFs were designed and used to carry out further local refinements and eliminate bad particles through 3D classifications. The resolutions of these local refined maps range from 3.6 to 4.4 Å.

A composite map was generated using the *volume maximum* command in ChimeraX (Pettersen, Goddard et al. 2021). DeepEMhancer was used to postprocess the maps (Sanchez-Garcia, Gomez-Blanco et al. 2021). Figures were made by ChimeraX.

### Protein domain identification

The symmetrical densities on C1 and C2 indicated in Fig. 1D were segmented and used in DomainFit (Gao, Tong et al. 2024) to identify the domains/proteins forming these densities (Fig. S2, Supplementary Table 2). The protein database comprises predicted models of 990 proteins identified in the mass spectrometry data of the WT axoneme (Kubo, Black et al. 2023). A total of 856 models were available and downloaded directly from the AlphaFold Database (https://alphafold.ebi.ac.uk/) (Jumper, Evans et al. 2021), and the remaining high-molecular-weight proteins were predicted by ColabFold (AlphaFold2) (Mirdita, Schutze et al. 2022).

The AlphaFold2-predicted models were divided into 1835 domains larger than 30 amino acids on the basis of the Predicted Alignment Error scores (Gao, Tong et al. 2024). These domains were fitted into each density filtered at 4 Å using 400 initial search position placements. For all the segmented densities, the P values of the top hits were significant, which allowed us to confidently assign the domain types to the density (Supplementary Table 2).

The top hits for each density were carefully reviewed and compared to crosslinks found in the crosslinking/mass spectrometry study of *Tetrahymena* cilia (McCafferty, Papoulas et al. 2023). In addition, the emPAI score of each protein was determined to ensure that the candidates had relatively similar abundances. The size of the protein was also used to filter out proteins that were too large, considering the surrounding densities. A summary of the identified proteins and the rationale for their assignment is provided in Supplementary Table 1.

For Tubulin Surface Density 2, the domain determined was a DPY-30 motif, which normally forms a dimer. AlphaFold2 was used to predict all the homodimers of proteins in the ciliome with the DPY-30 motif, and DomainFit was run with these dimers (Fig. S1F). Owing to the simple organization of the DPY-30 motif, the correct DPY-30 motif-containing protein could not be identified. I7LWQ5 was chosen for *in vivo* localization experiments because it crosslinked to the top hits of other densities.

There are still densities to which domains are not assigned, such as part of the Spike density. We were not able to identify the bridge proteins or the helical bundles above C2 confidently because of the flexible nature of the tip CP. All identified proteins repeat every 8 nm. It is likely that some of the proteins we could not identify have repeats different from 8 nm.

### Modelling

On the basis of the identified domain, we modelled the rest of the proteins by either predicting the model of the full-length protein using AlphaFold2 (Jumper, Evans et al. 2021) or by predicting a complex of the proteins of interest with a tubulin dimer using AF_unmasked (Mirabello, Wallner et al. 2023). We manually adjusted the model in Coot (Emsley, Lohkamp et al. 2010) and performed real space refinement using Phenix (Afonine, Poon et al. 2018).

### Sequence analysis

All amino acid sequences of the proteins used in this study were obtained from the UniProt database. TtCFAP213 and CrFAP213 were first aligned, and the alignment was converted into an HMM profile via HMM version 3.4 (Eddy 2011). The UniProt database was then searched with the HMM profile. A MEME search was run on the sequences to find a motif between 6 and 20 amino acids. The GYNPITGEY motif was identified with an E value of 3.5e-1550 (Bailey, Johnson et al. 2015). A MAST search of the identified motif was then run on the ciliome of *Tetrahymena* (Kubo, Black et al. 2023) and *Chlamydomonas* (Chlamyfp.org) (Pazour, Agrin et al. 2005). All sequence alignments were performed via Clustal Omega (Sievers, Wilm et al. 2011).

The sequence logo of the GYNPITGEY motif was created using https://skylign.org/.

### Gene editing

All the genes encoding the proteins listed in this study were edited via homologous DNA recombination, using a targeting plasmid containing the neo4 selectable marker. The gene segments necessary for targeting were amplified with primers 5F and 5R (which amplified a terminal region of the coding sequence) and primers 3F and 3R (which amplified a segment of the 3’ UTR). These amplified regions were subsequently cloned and inserted into the pNeo24-GFP plasmid (see Supplementary Table 3 for primer details). The resulting edited gene fragments were integrated into their native loci through biolistic bombardment of *Tetrahymena* cells, followed by paromomycin selection.

### Tetrahymena gene knock-in

To express TLP1, TLP2, CFAP213, C2D1, SPIKE2, and TTHERM_00579000 in fusion with a C-terminally positioned GFP tag under the control of the respective native promoters, approximately 1 kb fragments of the coding region without TGA and the 3’UTR of each gene were amplified using Phusion Hot Start II high-fidelity DNA polymerase (Thermo Fisher Scientific Baltics, Lithuania) and primers containing unique restriction sites at the 5’ and 3’ ends (Supplementary Table 3). After digestion with the restriction enzymes, the gene fragments were subsequently cloned and inserted into the pIFT52-GFP-neo4 plasmid digested with the same enzymes (Hazime, Zhou et al. 2021) to remove IFT52 gene fragments. Approximately 10 µg of the purified plasmids, pTLP1-GFP-neo4, pTLP2-GFP-neo4, pCFAP213-GFP-neo4, pC2D1-GFP-neo4, pSPIKE2-GFP-neo4, and TTHERM_00579000-GFP-neo4 each), were digested with MluI and XhoI restriction enzymes to separate the transgene from the plasmid backbone, precipitated onto 0.6 nm gold microcarriers (Bio-Rad #1652262) and introduced into CU428 *Tetrahymena* cells by biolistic transformation (Cassidy-Hanley, Bowen et al. 1997, Dave, Wloga et al. 2009). Transformed cells were selected on SPP supplemented with 100 µg/ml paromomycin and 1.5 µg/ml CdCl_2_. To promote transgene assortment, transformants were cultured on SPP media supplemented with increasing concentrations of paromomycin (up to 1 mg/ml) and decreasing concentrations of CdCl_2_ (up to 0.2 µg/ml).

### Superresolution - Structured illumination microscopy

For immunofluorescence, *Tetrahymena* cells were fixed and stained following the protocol described by Bre et al. (Bre, Redeker et al. 1996). The primary antibodies used included mouse monoclonal anti-polyglycylated tubulin AXO49 (1:200 dilution) (Bre, Redeker et al. 1996), anti-monoglycylated tubulin TAP952 antibody (1:5,000) (Bre, Redeker et al. 1998), and polyclonal anti-GFP antibodies (Rockland, 1:800). The secondary antibodies used were goat anti-mouse IgG-FITC and goat anti-rabbit-Cy3 (Jackson ImmunoResearch). Superresolution structured illumination microscopy (SR-SIM) imaging was performed using an ELYRA S1 microscope with a 63× NA 1.4 Oil Plan-Apochromat DIC objective. Optical slices were analysed using Fiji/ImageJ (Z projection tool) (Schindelin, Arganda-Carreras et al. 2012).

### Quantification

The length of the tagged tip CP proteins was quantified using the FilamentDetector version 0.4.7 plugin in FIJI (Mary and Rueden 2019).

### Cloning

The DNA sequence encoding human SPEF1 (UniProt ID: Q9Y4P9-1) cloned and inserted into pUC57 was ordered from Bio Basic Canada, Inc. The HsSPEF1 CH-domain (residues 1-125) was cloned via USER cloning (Bitinaite, Rubino et al. 2007) into a modified pHAT vector containing an N-terminal His tag, a C-terminal eGFP and a C-terminal StrepII tag (kind gift from G. Brouhard).

### Protein expression and purification

The resulting plasmids were subsequently transformed into *E. coli* BL21(DE3). Cultures were grown at 37°C until OD_600_ = 0.6 and then induced overnight with 0.5 mM IPTG at 18°C. The cells were subsequently resuspended in lysis buffer (50 mM Tris pH 7.0, 500 mM NaCl, 40 mM imidazole, 1 mM EGTA and 5 mM β-mercaptoethanol) supplemented with 2 μg/mL aprotinin (MilliporeSigma) and 100 μM leupeptin (MilliporeSigma) and were subsequently lysed via EmulsiFlex-C5 (Avestin). The lysate was cleared by centrifugation (1 hour, 50,000 × *g*) in a JA 25.50 rotor (Beckman Coulter), filtered and loaded onto a 5-mL HisTrap FF column (Cytiva). Proteins were eluted in elution buffer (lysis buffer with 300 mM imidazole). Fractions containing the protein of interest were incubated overnight in dialysis buffer (25 mM Tris, 300 mM KCl, 10 mM imidazole, 1 mM EGTA and 5 mM β-mercaptoethanol) with 3C protease and then loaded onto a HisTrap FF column (Cytiva). The eluted protein was then concentrated and loaded on a Superdex 200 10/300 GL (Cytiva) pre-equilibrated in size-exclusion chromatography buffer (20 mM Tris pH 7.5, 300 mM KCl, 1 mM EGTA and 1 mM dithiothreitol (DTT)).

The 3C protease was purified in-house using the pHN924 plasmid from Addgene.

### Cryo-ET preparation

*Bos taurus* purified tubulin (in-house) was incubated at 37°C for 1 hour in BRB80 supplemented with 1 mM DTT, 3 mM GTP and 20 μM paclitaxel (Millipore Sigma). Microtubules were centrifuged at 199,000 × *g* for 5 minutes to remove soluble tubulin and resuspended in BRB80. Five micromolar Taxol-stabilized microtubules were then incubated at an equimolar ratio with HsSPEF1-CH for 30 minutes with 5 nm gold beads (Cytodiagnostics, 1:3 ratio). Three microliters of sample were applied to negatively glow-discharged (10 mA, 10 s) C-Flat Holey thick carbon grids (Electron Microscopy Services #CFT312-100) for 1 min inside a Vitrobot Mk IV (Thermo Fisher Scientific). The chamber was set to 23°C with 90% humidity. The grids were blotted for 4 s at a blot force of 15 and plunged into liquid ethane.

### Tilt series acquisition

Tilt series were acquired on a Titan Krios electron microscope (Thermo Fisher Scientific) operated at 300 kV and equipped with a BioQuantum energy filter (Gatan, Inc.) and a K3 Summit (Gatan, Inc.) electron detector. Tilt series were collected using SerialEM (Mastronarde 2005) at 42,000× magnification using a grouped dose□symmetric scheme from −60° to 60° with an increment of 3°. The defocus for each tilt series ranged from −2 to −4 μm. The total dose for each tilt series was 164 e^−^ per Å^2^. For each view, a movie of 10 frames was collected. The pixel size at 42,000× was 2.12 Å. Frame alignment was performed with Alignframes (Mastronarde and Held 2017). Tomograms were reconstructed by IMOD (Kremer, Mastronarde et al. 1996). The aligned tilt series were manually inspected for quality and sufficient fiducial presence. Batch reconstruction was performed in IMOD. The tomograms shown in the figures were denoised using the SIRT-like filter provided in IMOD.

### Subtomogram averaging

CTF estimation of the tomogram was performed with WARP (Tegunov and Cramer 2019). The subtomograms were generated through a set of Dynamo (Castano-Diez, Kudryashev et al. 2012) scripts for microtubule alignment (https://github.com/builab/dynamoMT). In brief, microtubules were picked using IMOD by tracing lines along their centre. The 8-nm overlapping subtomogram centers were interpolated along the microtubule tracing lines. All subtomograms belonging to the same microtubules were roughly aligned and averaged to generate a microtubule average. Using the microtubule averages, we could align and classify them with the 13-PF and 14-PF references. After that, the 13-PF and 14-PF microtubules were separated, and subtomogram averaging was performed independently on each set of subtomograms for the 13- and 14-PF microtubules. The subtomogram coordinates and orientations were then exported to Relion 5 (Burt, Toader et al. 2024) for refinement. Finally, 2571 and 2831 subtomograms were used for the average maps of 13 and 14-PF microtubules, resulting in resolutions of 19 and 17 Å, respectively.

For microtubule pair subtomogram averaging, the picking of microtubule pairs was performed using IMOD by tracing lines along the center of the pair. The subtomograms were then picked at 8-nm intervals (approximately 100 particles per microtubule pair) along the traced lines and averaged independently.

For visualization, we transformed the subtomogram averages into the original tomogram coordinates of each subtomogram in ChimeraX via the subtomo2Chimera script (Bui 2022).

Models of 13-PF and 14-PF microtubules with HsSPEF1-CH binding at the seam was built on the basis of the 13-PF and 14-PF microtubule PDB models 5SYC (Zhang, Alushin et al. 2015, Kellogg, Hejab et al. 2017) and 3JAT (Zhang et al., 2015). The AlphaFold2 model of HsSPEF1-CH was placed at the seam of the 13-PF and 14-PF microtubule model based on the structure of TtSPEF1A binding at the seam of the C2 microtubule.

### TIRF microscopy

The microscope setup and the microtubule dynamics assays were performed as described previously (Wieczorek, Bechstedt et al. 2015). In brief, GMPCPP-stabilized microtubule seeds were introduced into a flow cell and immobilized onto a silanized cover glass via antibodies. TIRF images were acquired with sequential excitation with a 491 nm laser (HsSPEF1-CH channel) and a 561 nm laser (AF546-labelled microtubule channel) at intervals of 10 s/frame for 180 frames. A polymerization mixture containing 8 μM tubulin (labelled with 10% AF546) or 8 μM tubulin and 100/400/750/1000/2000 nM HsSPEF1 was introduced into the flow cell between the 1st and 2nd frames during image acquisition. The polymerization buffer was BRB80 supplemented with 1 mM GTP, 0.1 mg/ml BSA, 0.2 mg/ml k-casein, 29 mM KCl, 10 mM DTT, 1x oxygen scavenging cocktail (250 nM glucose oxidase, 64 nM catalase, 40 mM D-glucose), and 0.01% methylcellulose. The assays were performed at 35°C.

The analysis of microtubule dynamics was conducted using a combination of manual line fitting and customized Python scripts. In brief, kymographs of dynamic microtubules were generated via ImageJ. Manual line fitting over the first triangle of a given kymograph, which represents the first growth-shrinkage event of the microtubule over the movie duration, was conducted to generate xy coordinates that contain the temporal information of the dynamic microtubules. The xy coordinates were then used to calculate dynamic parameters, including the microtubule length, lifetime, growth rate, time to nucleate, and shrinkage rate. Microtubule rescue was defined as true if the microtubule was more than 5 pixels from the seed upon rescue. The rescue probability was then calculated by dividing the number of true rescue events by the total number of events for any given replicate.

### Statistics

Statistical analyses were carried out using GraphPad Prism version 10.3.0.

## Acknowledgement

We thank all the members of the Bui laboratory for their critical reading of the manuscript. We thank Drs. Mike Strauss, Kaustuv Basu, and Kelly Sears (Facility for Electron Microscopy Research at McGill University) for helping with data collection. KHB is supported by grants from the Canadian Institutes of Health Research (PJT-156354) and the Natural Sciences and Engineering Research Council of Canada (RGPIN-2022-04774). JG is supported by National Institutes of Health grant R01GM135444. TL is supported by the Funds de Recherche du Québec Sante (FRQS) fellowship (353841).

## Author contributions

Conceptualization: KHB. Methodology: T.L. Formal analysis: TL, EW, MP, AT, EJ, WP, CB, MV-P, DW, and KHB. Resources: JG, GB, DW and KHB. Writing–original draft: TL and KHB. Writing–review & editing: TL, KHB, JG, EJ, and DW.

## Data and code availability

○ The structural data (atomic coordinates and structural factors) have been deposited in the Protein Data Bank (https://www.rcsb.org) and EMDB (https://www.ebi.ac.uk/emdb/). Single-particle map of the tips of CP EMD-XXXX and PDB YYYY. Subtomogram average map of the microtubule-hsSPEF1-CM structure: EMD-ZZZZ.
○ The code for subtomogram alignment and averaging of the microtubules is available at https://github.com/builab/dynamoMT.

## Supplementary material

**Supplementary Movie 1:** Visualization of the single-particle cryo-EM tip-CP map and structure from a tomogram slice of a *Tetrahymena* cilium (from Legal et al., 2023). The colours in the map and structure represent different tip proteins and are consistent with the colours in Figure 1.

**Supplementary Movie 2:** Visualization of a tomogram of microtubules decorated with recombinant HsSPEF1-CH. The subtomogram averages are then overlaid on the tomogram for clear visualization of the HsSPEF1-CH density (green).

**Supplementary Table 1:** Identified proteins in the tip CP along with the residues modelled, the emPAI score and the rationale for their assignments.

Table S1.xlsx

**Supplementary Table 2:** Top 10 hits from DomainFit for each density shown in Fig. S1E. The identified proteins are highlighted in yellow.

Table S2.xlsx

**Supplementary Table 3:** Primers used for GFP tagging of tip CP proteins

Table S3.xlsx

